# EDA Fibronectin Microarchitecture and YAP Translocation During Wound Closure

**DOI:** 10.1101/2024.09.23.614581

**Authors:** Jennifer Patten, Patrick Halligan, Ghazal Bashiri, Michael Kegel, Jacob D Bonadio, Karin Wang

**Affiliations:** Department of Bioengineering, Temple University, Pennsylvania

## Abstract

Fibronectin (Fn) is an extracellular matrix glycoprotein with mechanosensitive structure-function. EDA Fn, a Fn isoform, is not present in adult tissue but is required for tissue repair. Curiously, EDA Fn is linked to both regenerative and fibrotic tissue repair. Given that Fn mechanoregulates cell behavior, Fn EDA organization during wound closure might play a role in mediating these differing responses. One mechanism by which cells sense and respond to their microenvironment is by activating a transcriptional co-activator, Yes-associated protein (YAP). Interestingly, YAP activity is not only required for wound closure, but similarly linked to both regenerative and fibrotic repair. Therefore, this study aims to evaluate how, during normal and fibrotic wound closure, EDA Fn organization might modulate YAP translocation by culturing human dermal fibroblasts on polydimethylsiloxane (PDMS) substrates mimicking normal (soft: 18 kPa) and fibrotic (stiff: 146 kPa) wounded skin. On stiffer substrates mimicking fibrotic wounds, fibroblasts assembled an aligned EDA Fn matrix comprising thinner fibers, suggesting increased microenvironmental tension. To evaluate if cell binding to the EDA domain of Fn was essential to overall matrix organization, fibroblasts were treated with Irigenin, which inhibits binding to the EDA domain within Fn. Blocking adhesion to EDA led to randomly organized EDA Fn matrices with thicker fibers, suggesting reduced microenvironmental tension even during fibrotic wound closure. To evaluate if YAP signaling plays a role in EDA Fn organization, fibroblasts were treated with CA3, which suppresses YAP activity in a dose-dependent manner. Treatment with CA3 also led to randomly organized EDA Fn matrices with thicker fibers, suggesting a potential connected mechanism of reducing tension during fibrotic wound closure. Next, YAP activity was assessed to evaluate the impact of EDA Fn organization. Interestingly, fibroblasts migrating on softer substrates mimicking normal wounds increased YAP activity but on stiffer substrates, decreased YAP activity. When fibroblasts on stiffer substrates were treated with Irigenin or CA3, fibroblasts increased YAP activity. These results suggest there may be disrupted signaling between EDA Fn organization and YAP translocation during fibrotic wound closure that could be restored when reestablishing normal EDA Fn matrix organization to instead drive regenerative wound repair.

## Introduction

Wound healing in adult skin lacks the ability to recapitulate uninjured tissue structure-function, forming instead fibrotic tissue with limited functionality. Regeneration restores wounded tissue to its original organization, comprising controlled deposition and assembly of extracellular matrices (ECM) with basket-weave pattern or random organization^1^ through poorly understood or unknown mechanisms. Inducing regeneration is the holy grail of wound healing. Repair, the most common result of wound healing, often leads to increased deposition and dysregulated ECM assembly with excessively aligned fibers resulting in fibrotic tissue.^2,3^ The ECM^4^ regulates migration, morphology and fibrosis^5–7^ and is a unique target for therapeutic control of the fibrotic response.^8–10^

Mechanical forces, including tissue stiffness, play a key role in determining wound fate, with mitigation of mechanotransduction-based signaling decreasing the fibrotic matrix response.^11–14^ Among the ECM proteins, fibronectin (Fn) distinguishes itself through its mechanosensitivity.^15,16^ FnIII modules within Fn lack disulfide bonds and therefore make it mechanically malleable, allowing the fibronectin fiber to elongate up to 4x its resting length, exposing or disrupting various binding domains.^17^ Fn is a ∼500 kDa dimeric matrix glycoprotein that contains a plethora of binding domains for cell adhesion, growth factor immobilization, and matrix protein binding.^16,18^ Through these binding sites, fibronectin can variably dictate many cellular functions, including migration, adhesion, differentiation, and proliferation, all essential wound healing functions. Therefore, it is unsurprising that Fn drives cell behavior throughout all phases of wound healing.^19–23^

During clotting, circulating plasma Fn helps stem blood flow and begin provisional ECM formation at the wound site.^20,24^ During inflammation, cells begin secreting cellular derived Fn that will then be replaced by a more permanent collagen matrix in the proliferation phase.^25–27^ During the remodeling phase, the Fn matrix either returns to a normal homeostatic response,^20,21^ or a chronic increase in Fn deposition and matrix alignment leads to an excessive fibrotic response.^28^ Fibrosis is characterized by a chronic over-alignment of the matrix associated with increased tissue stiffness; however, too disorganized and loose a Fn matrix during wound healing will not promote wound closure and will lead to worse wound outcomes.^29–31^

One way that Fn controls cellular behavior is through the expression of different isoforms; depending on which exons are included, Fn’s isoform structure-function changes.^18,25,26,32,33^ The Extra Domain A (EDA) region is a cell-derived Fn isoform normally associated with fetal development and is only expressed in adults during wound healing and pathogenesis.^32,34^ The EDA fibronectin’s exon is located between FnIII^11–12^ at a central hub of integrin binding and growth factor sites and is suggested to be alternatively spliced at the wound site in response to microenvironmental cues,^3^ including increased mechanical tension.^35^ EDA, an extra FnIII module inserted into Fn, alters 3D conformation of its fibers,^36^ can bind additional integrins,^3^ and is heavily involved in inflammatory signaling.^37–42^ The EDA Fn exon is essential for wound closure ^21,43^ and promotes regeneration.^44–46^ However, the chronic overexpression of EDA Fn at the wound site is directly correlated to the strength of the fibrotic response.^47–50^ How EDA Fn regulates these disparate wound outcomes remains unclear; the altered mechanical profile of EDA Fn compared to fibronectin isoforms lacking the EDA domain is likely to hold some answers.

Fn, as the provisional matrix, bears the primary mechanical load before being supported and replaced by the permanent collagen matrix as it is deposited.^27,51,52^ EDA Fn is a cryptic binding site conformationally blocked at equilibrium; it requires unfolding to unlock its binding domain, such as through increased mechanical tension.^41,53^ And mechanically sensitive EDA Fn drives a variety of cellular functions including migration, fibrosis, and differentiation. EDA Fn likely assumes more mechanical loading than EDA negative Fn to unlock its cryptic domain.^41^ Increased ECM tension also increases EDA Fn expression.^35^ As such, it is likely that EDA Fn bearing more load would expose the cryptic EDA domain to increase EDA-specific cell binding to alter wound healing responses. Therefore, controlling ECM microarchitecture is a potential mechanotransduction-based mechanism to restore normal cell behavior during wound healing. It is likely that increased mechanical tension is an important regulator of normal or fibrotic wound fate decided by EDA Fn. Therefore, this study seeks to investigate how substrate stiffness, mimicking either normal or fibrotic wounds, changes EDA Fn matrix microarchitecture during wound closure.

Furthermore, the mechanism by which EDA Fn matrix microarchitecture mechanoregulates cell signaling during wound closure remains unclear. Yes associated protein (YAP) is a mechanotransduction-based transcriptional coregulator^54^ that is known to be activated by Fn^53,54^ and mechanical stresses^55^ (*i.e*., substrate stiffness).^56^ Like EDA Fn, YAP is required for wound closure^57^ and linked to regenerative tissue repair.^45,58,59^ Again similarly to EDA Fn, excessive YAP signaling also drives a fibrotic response over time.^60^ YAP is sensitive to viscoelastic mechanical stimuli over elastic input, and activates various mechanosensing cell signaling functions beyond migration and fibrosis, including wound healing behaviors such as cell attachment/spreading,^61^ proliferation,^53^ and apoptosis.^62^ Why YAP can successfully drive such a variety of contradictory cell behaviors may be due to its redundancy as a mechanotransduction signaling effector upregulating target genes in response to many different signaling pathways.^6,13,63–67^ However nuanced mechanical influences (*i.e.*, matrix-mediated signaling, structural cues) on the mechanoregulator YAP will likely will provide insight into the dichotomy of YAP signaling.^68–70^ Given that EDA Fn and YAP demonstrate similar contradictory functions dictating either normal or fibrotic wound repair and are both mechanically sensitive, we hypothesized that YAP is a mechanotransduction signaling effector potentially controlled by EDA Fn during wound closure. To address our fundamental question of whether EDA Fn and YAP are on the same signaling axis to control wound fate, we developed a 2D wound closure assay to investigate if substrate stiffnesses mimicking normal or fibrotic wounds would similarly control EDA Fn matrix microarchitecture and YAP signaling.

Fn is a mechanically sensitive ECM protein with an isoform, EDA Fn, whose expression within wounds variably drives normal or fibrotic healing. Though EDA Fn is found in wounded and fibrotic adult tissue and is correlated with yes-associated protein (YAP) nuclear translocation, the mechanisms by which EDA Fn matrix microarchitecture mechano-regulates YAP nuclear translocation to drive normal or fibrotic healing remains unclear. Therefore, by controlling substrate stiffness to mimic either normal or fibrotic wounds and inhibiting EDA-specific cell binding to Fn or attenuating Yes-associated protein (YAP) signaling, we seek to evaluate if differential EDA Fn microarchitecture mediates YAP activity.

## Experimental Setup

### Cell culture

Human dermal fibroblasts, adult (HDFa) cells (PCS-201-012, ATCC) were cultured in DMEM 1X, 10% FBS (high serum) and 1% P/S. Cells passage 8 or younger were used in this assay. Cells were seeded at 45,000 cells/well in a 12 well plate 24 hours pre-wounding. Cell culture area was constrained by a 100 mm^2^ PDMS O-ring to create a cell seeding density of 45,000 cells/cm^2^.

### Material synthesis

Polydimethysiloxane (PDMS) substrates were crosslinked at 49:1 and 40:1 ratios of base:crosslinker (Sylgard 184, Ellsworth, 50-366-794) to produce varying elastic moduli at 18 kPa and 146 kPa, respectively. The PDMS was weighed into each well of a 12 well plate, 0.25 g to create a thin layer ∼500 µm for culturing the cells. PDMS was cured for 4 hours at 60°C to complete the curing process. Substrates were washed with phosphate buffered saline (PBS) for 2 hours (2×30 minutes and 1xhour incubations) to clear the substrate surface. Plates were sterilized with ultraviolet light for 15 minutes. Masks, to simulate wound area, were cut to be 0.5 mm width from 10:1 crosslinked PDMS cured at 60°C for 4 hours. PDMS masks and O-rings were sterilized per face for 15 minutes each. Plasma fibronectin (33016015, ThermoFisher) at a concentration of 30 µg/mL was used to coat (1 hour incubation, 24°C) the PDMS substrate surface pre-mask placement to enable cell attachment. After plasma fibronectin attachment, the masks were placed in the center of each well. HDFa cells were seeded around each mask and left to adhere for 30 minutes at 37°C and 5% CO2 before a media wash step was performed. Cells were then cultured to confluency around the masks for 24 hours before mask removal. To simulate wound, masks were removed, and fresh media was added to each well at the assay start.

### Mechanical testing

Compression testing was performed on the BOSE Electroforce 3230, where PDMS samples were compressed to 10% of their sample thickness at a rate of 0.0166 mm/s. Stress was found by the equation: σ=Force/Area, and strain was calculated by ɛ=ΔL/L, displacement over total sample length. The linear region of the resulting stress-strain plots was found by linear regression with an R^2^≥0.99. The slope of the linear regression was taken as the Young’s modulus of the PDMS.

### Conditional groups and controls

Media was changed every 24 hours during the wound assay. The inhibition conditions were Irigenin, 50 µM (PHL83512-10MG, Millipore Sigma),^71^ and YAP inhibitor CA3^62^ (CIL56, S8661, SelleckChem) at 0.5 µM concentrations. Each inhibitor was resuspended in dimethylsulfoxide (DMSO, 31 762 5ML, FisherScientific), and a DMSO wound assay control condition was run at 45 µL/mL, corresponding to the larger inhibitor volume added to reach the desired concentration.

### Assay endpoint and immunostaining

Plates were fixed for one hour in 4% paraformaldehyde at 4, 24, 48, or 72h time points. Immunostaining was performed with phalloidin (1:250, f-actin cytoskeleton, A12380, ThermoFisher), DAPI (1:5000, D1306, ThermoFisher), IST-9 (1:400, EDA Fn, Abcam), AlexaFluor647 (1:100, secondary antibody for EDA Fn, A32728 ThermoFisher), Fn antibody (1:100, PA5-29578, ThermoFisher), YAP antibody (1:200, D8H1X, 14074S, Cell Signaling Technology), and AlexaFluor 488 (1:100, secondary antibody for Fn and YAP, A-11008, ThermoFisher).

### Confocal imaging and analysis

Confocal images were obtained from within each wound using an Olympus Fluoview FV1200, 30x oil objective, 2 µm thickness step slices, 0.75 µm pinhole. Lasers and laser powers used were the 405 at 4%, HV 507 (DAPI), 488 at 10%, HV 400 (AlexaFluor488), 543 at 21%, HV 600 (phalloidin), 635 at 3%, HV 440 (AlexaFluor 647). The Olympus confocal microscope outputs OIF files with the individual laser and brightfield channels embedded into one image. Images were processed in ImageJ Fiji to project the max intensity of the z-slices taken into a single image according to the following preprocessing pipeline: the channels were split, separating out 5 images, corresponding to the 4 lasers and brightfield. The maximum intensity of the z-slices was then projected to a new maximum intensity grayscale image for all laser channels, excluding brightfield. Max intensity images were saved for further analysis and then merged into a colored composite. These max intensity images were processed through TWOMBLI, an ImageJ Fiji macro which creates a binary mask from the fluorescent ECM fibers to analyze its microarchitecture.^72^

Further matrix microarchitecture analysis upon these EDA Fn confocal images were performed to evaluate fiber width and fiber angle using CT-FIRE, a separate matrix analysis software that identifies individual fiber metrics within a matrix via utilization of the curvelet transform and denoising techniques.^73^ The CT-FIRE input parameters of minimum fiber width, length were adjusted to fit the image resolution, and the CT-FIRE analysis was performed. Image exclusion criteria included visible deformations to the PDMS substrate.

In ImageJ Fiji, max intensity projected images of DAPI, YAP, and actin were used to visualize the cell landscape and obtain accurate nucleic and cytoplasmic YAP fluorescent intensity per cell. Each measurement recorded the area, minimum and maximum gray value, integrated density, and mean gray value of the outlined region. Cytoplasmic and nuclei area and associated mean fluorescent intensities were found using the ImageJ Fiji intensity density function (mean gray intensity multiplied by area) and normalized for area to calculate the nucleic/cytoplasmic YAP ratios from the relative fluorescent intensities. Measurements of ∼20 cells were taken per image, ≥5 images per condition, save for the 4h condition which had N≥18. Inclusion criteria for cell measurement included <90% cell volume in field of view and distinct cell boundaries.

### Statistical analysis and Data Processing

GraphPad Prism 9 was used to run one-way ANOVA with Tukey’s post hoc analysis. Outliers were defined as outside of two standard deviations from the mean of each condition and were removed. As TWOMBLI alignment, branching, density, and CT-FIRE fiber width and angle were sourced from the same images, these parameters were treated as paired for outlier removal from these data sets. Similarly, the YAP nuclear/cytoplasmic analysis and whole cell area analysis were also treated as paired for outlier removal. Data are presented graphically as box and whisker plots in main figures and tables of mean and standard deviations in supplemental; for clarity, in-text citations show means with standard deviations. Schematics were created in Biorender.

## Results & Discussion

### 2D biomimetic skin wound assay

Although *in vivo* studies are the gold standard for wound studies,^74^ *in vitro* models such as the wound closure assay are recognized as important alternative methods to investigate cell-matrix interactions and dynamics.^75^ This 2D wound closure assay investigates EDA Fn matrix assembly and YAP nuclear translocation on two substrate stiffnesses, 18 kPa^76,77^ and 146 kPa,^78^ representing wounded and fibrotic human skin stiffnesses, respectively.

Polydimethylsiloxane (PDMS) was selected as the substrate because it demonstrates viscoelastic properties similar to human skin.^32,37^ By altering the base:crosslinking ratio of PDMS, the stiffness can be selectively varied. First, compression testing (Fig. 1A) was performed on various PDMS fabrications (Fig. S1) to select the best base:crosslinking ratio that would mimic wounded and fibrotic skin stiffnesses (Fig. 1B). From the compression testing, the Young’s moduli were found to be 18 kPa and 146 kPa for the 49:1 and 40:1 crosslinking ratios cured at 60°C in the oven, respectively. Plasma fibronectin was adsorbed onto the hydrophobic PDMS substrates to facilitate cell adhesion and to provide the plasma fibronectin that is the first of the fibronectin isoforms to be recruited to the wound site during the clotting stage.^20^ Adult human dermal fibroblasts, HDFas, were cultured to confluency around a PDMS mask; removal of the mask simulated wounding and the beginning of the wound assay (Fig. 1C). The fibroblast assembled fibronectin matrices post-wounding (mask removal) were immunostained and evaluated by confocal imaging (Fig. 1D). Matrix organization analyses were performed using two matrix analysis software, the ImageJ macro TWOMBLI to extract alignment, density, branching, and the Matlab-based CT-FIRE to extract fiber width and orientation (Fig. 1E). Tracing YAP in nuclear and cytoplasmic compartments was run concurrently to investigate a potential EDA-YAP signaling axis during wound closure (Fig. 1F).

**Figure 1:**
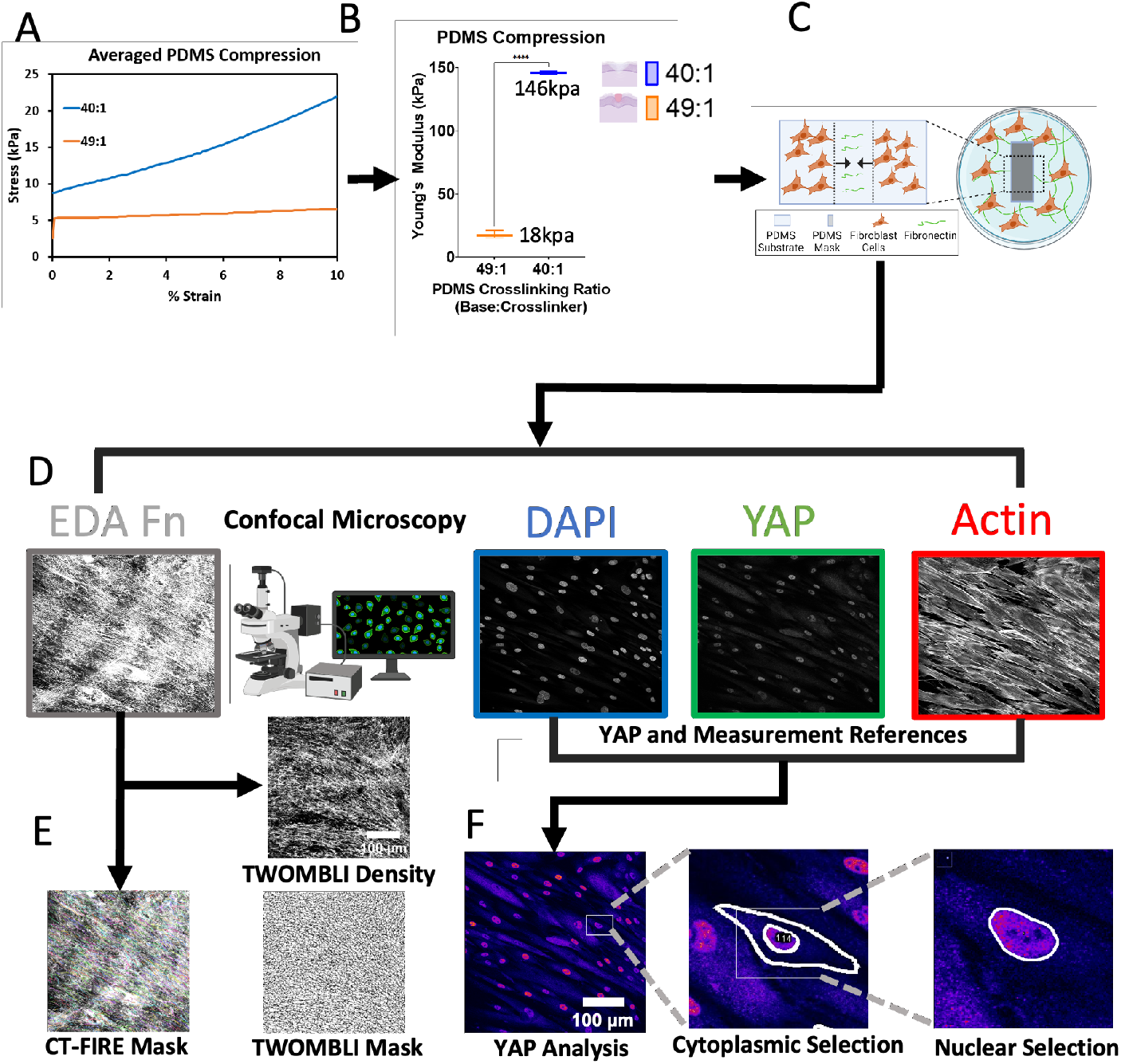
Fabricating PDMS substrates for wound assays and image analysis pipeline. **A**. Compression testing to 10% of material thickness was performed on polydimethylsiloxane (PDMS) substrates at varying base:crosslinker ratios to verify target stiffnesses. **B**. Young’s Moduli, the linear portion for each stiffness, was calculated by applying a linear regression fit with an R^2^≥0.99. One-way ANOVA, Tukey’s posthoc analysis, ****P<0.0001. **C**. The wound assay was created by adsorbing plasma fibronectin onto the PDMS substrates, placing a PDMS mask, then culturing adult human dermal fibroblasts, HDFa, around the mask until confluency. Mask removal simulated beginning of wound assay at 0h. **D**. Confocal microscopy was used to image fluorescently tagged EDA Fn, actin, DAPI, and YAP. **E.** Matrix microarchitecture image analysis pathway. CT-FIRE and TWOMBLI analysis was run from the processed EDA Fn images (max intensity across the Z-stack projected into a single image). TWOMBLI applied a high-density matrix (HDM) threshold from which TWOMBLI generated a binary mask with a matrix of dark lines depicting detected matrix and white space marking any elements that did not meet the HDM threshold applied by TWOMBLI. TWOMBLI outputs include matrix architecture quantifications of alignment, density, and number of branch points. A similar matrix analysis algorithm, CT-FIRE, was also run on the processed EDA FN images to identify fiber width and angle. **F**. DAPI, actin, and YAP images were used to extract YAP nuclear and cytoplasmic measurements. YAP image was converted from grayscale to a preset “fire” color scheme, to clear cell outlines without changing the pixel/fluorescence data. From left to right shows the cell selection (left), cytoplasmic (middle) and nucleic (right) YAP freehand outlining process from which the fluorescent intensity was obtained via ImageJ Fiji measurement. Schematics were created in Biorender.

### EDA Fn matrix microarchitecture is time- and substrate stiffness-dependent

First, human dermal fibroblasts migrating into wounds on PDMS substrates mimicking normal (soft: 18kPa) and fibrotic (stiff: 146 kPa) wounded tissue were evaluated for the presence and timing of EDA fibronectin matrix assembly (Fig. 2A). On both PDMS substrates mimicking normal and fibrotic wounds, EDA Fn matrices were assembled within wounds as early as 24h. Noticeably, these matrices when stained with a polyclonal antibody recognizing a region within the N-terminus of Fn (PA5-29578, ThermoFisher; binds amino acids between 396 and 689 on fibronectin, comprising parts of the FNII1-2FNI7-9FNIII1 domains), revealed only the plasma Fn coating adsorbed onto the PDMS substrate to facilitate cell adhesion and plasma Fn matrix organization at later time points (Fig. S2). Therefore, within both our soft (normal) and stiff (fibrotic) PDMS-based wound closure assay, HDFa deposited EDA Fn dictates much of the early matrix assembly.

**Figure 2:**
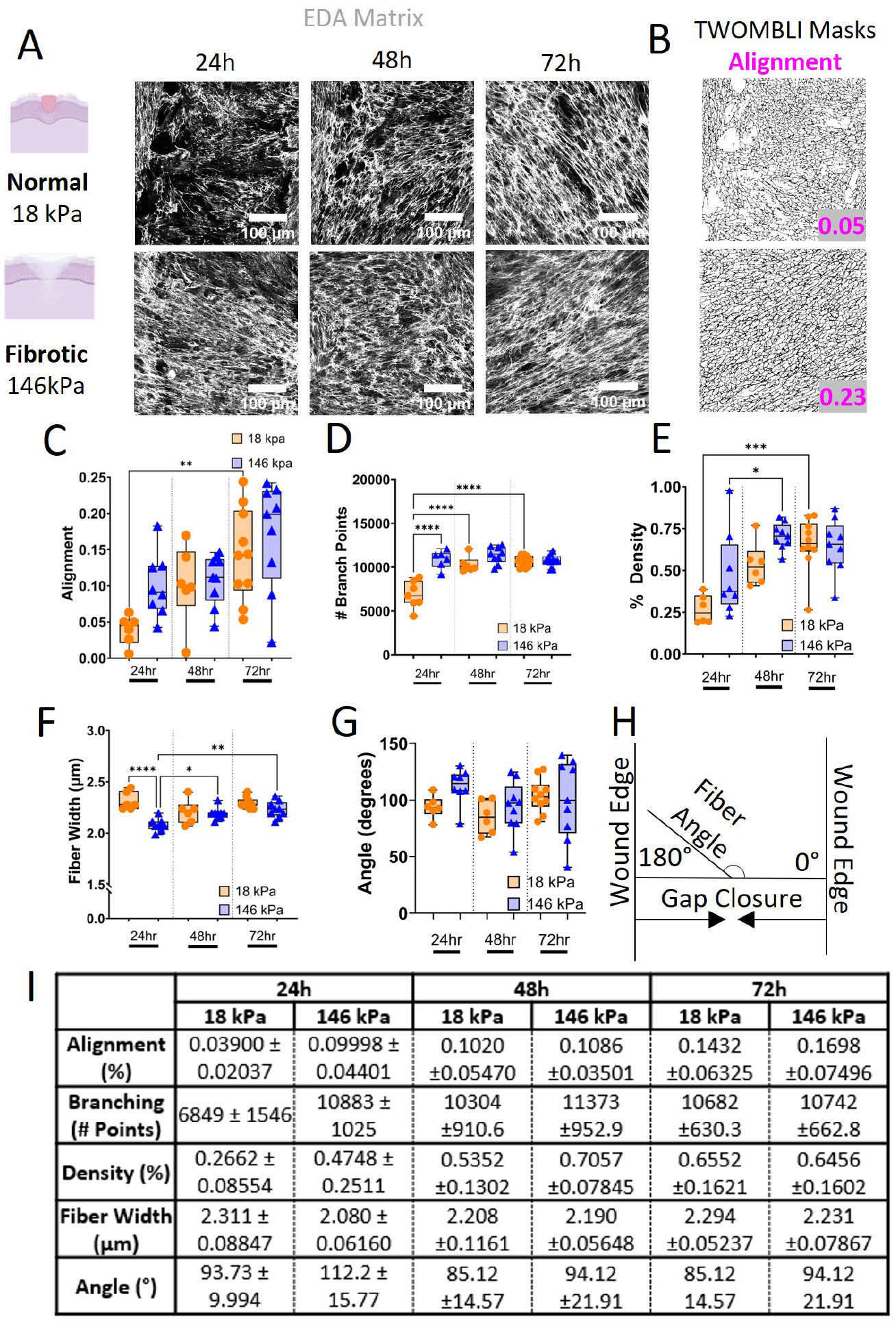
EDA fibronectin microarchitecture post-wounding. **A.** Fluorescently labeled EDA Fn images were taken by confocal microscopy, scale bars 100 µm. **B**. Example matrix masks and alignment values generated from our samples by the ImageJ macro, TWOMBLI. From top to bottom, masks shown correspond to images shown in **A**: 24h, 18 kPa and 72h, 146 kPa. **C**. EDA fibronectin matrix alignment returned by TWOMBI for each condition, **P=0.0066. **D**. Number of EDA fibronectin matrix branch points as calculated by TWOMBLI. ****P<0.0001. **E**. EDA Fn matrix density as found by TWOMBLI analysis. *P=0.0482, ***P=0.0003. **F**. CT-FIRE analysis of fiber width, µm. *P=0.046, **P=0.002, ****P<0.0001. **G**. CT-FIRE analysis of fiber angle, **H**. measured in degrees from horizontal (0 180). All N≥5, one-way ANOVA, Tukey’s post-hoc analysis. **I**. Table of means ± standard deviations for the matrix analysis metrics, alignment, branching, density, fiber width, and fiber angle.

To quantify EDA Fn matrix microarchitecture, confocal images (Fig. 2A) were analyzed by an ImageJ Fiji macro, TWOMBLI, which quantified matrix alignment, branching, and density by generating a mask based on the signal obtained from the fluorescently labeled EDA Fn (Fig. 2B).^72^

EDA Fn matrices assembled by fibroblasts migrating into wounds reveal distinct matrix architectural changes across time and substrate stiffnesses (Fig. 2). Fibroblasts on stiffer substrates mimicking fibrotic wounds assembled EDA Fn matrices that were more aligned (146 kPa: 24h, 0.09998±0.04401; 48h, 0.1086± 0.03501; 72h, 0.1698±0.070496) than EDA Fn matrices assembled by fibroblasts on softer substrates mimicking normal wounds (18 kPa: 24h, 0.039±0.02037; 48h, 0.102±0.0547; 72h, 0.1432±0.06325) (Fig. 2C,I). Matrix fibers are randomly organized in unwounded skin. During wound repair, a certain degree of matrix alignment is necessary to facilitate wound closure. However, prolonged and excessive matrix alignment drives an increased fibrotic response.^29–31^ Similarly, we demonstrate that EDA Fn is more aligned over time in fibrotic wounds.^79^

We also evaluated EDA Fn branching or number of branch points (Fig. 2D,I). When used in conjunction with density, branching can be used as an indication of matrix porosity.^72^ We see that fibroblasts on stiffer substrates mimicking fibrotic wounds assembled more branched EDA Fn matrices (146 kPa: 24h, 10883±1025; 48h, 11373±952.9; 72h, 10742±662.8) than that assembled by fibroblasts on softer substrates mimicking normal wounds (18 kPa: 24h, 6849±1546; 48h, 10304±910.6; 72h, 10682±630.3). Matrix density was quantified by the fluorescent matrix signal detected in each field of view, or the amount of matrix assembled at each time point.^72^ Analysis of matrix density (Fig. 2E,I) demonstrates that fibroblasts migrating on stiff substrates mimicking fibrotic wounds (146 kPa: 24h, 0.4748±0.2511; 48h, 0.7057±0.07845; 72h, 0.6456±0.1602) overall assembled more EDA Fn than that of fibroblasts migrating on soft substrates mimicking normal wounds (18 kPa: 24h, 0.2662±0.08554; 48h, 0.5352±0.1302; 72h, 0.6552±0.1621). EDA

Fn branching and matrix density both plateau over time, suggesting that there may be a threshold to the density of EDA Fn synthesized, deposited, and assembled within wounds. Matrices undergo remodeled after assembly.^80^ The half-life of Fn is 24-72h,^81–83^ indicating that Fn matrices are turning over in order to maintain tissue functionality. Moreover, Fn matrix remodeling is responsive to cell and microenvironmental forces.^8,84,85^ Our data suggests that there needs to be sufficient matrix assembled within a wound, prior to matrix remodeling.

Next, individual EDA Fn fiber width was measured through CT-FIRE (Fig 2F,I). EDA Fn fibers assembled by fibroblasts migrating into wounds on stiffer PDMS substrates mimicking fibrotic wounds (146 kPa: 24h, 2.080±0.0616; 48h, 2.190±0.05648; 72h, 2.231±0.07867) were thinner than that of EDA Fn fibers assembled by fibroblasts migrating into wounds on softer PDMS substrates mimicking normal wounds (18 kPA: 24h, 2.311±0.08847; 48h, 2.208±0.1161; 72h, 2.294±0.05237). The initially thinner EDA Fn fibers suggest that the aligned EDA Fn matrix is under high amounts of tension. And at later timepoints, EDA Fn fibers assembled by fibroblasts on fibrotic substrates gradually became thicker, suggesting that there is less mechanical load so that EDA Fn fibers may relax. It is likely that the provisional EDA Fn matrix may be on its way to being remodeled into a collagen matrix,^27^ therefore thicker fiber widths could be due to the presence of collagen in the ECM, shielding the fibrotic (146 kPa) EDA Fn fibers from bearing a majority of the mechanical load. We also quantified EDA Fn fiber angles (Fig. 2G,I) against the horizontal plane (Fig. 2H). The horizontal origin plane was set parallel to the direction fibroblasts must travel to connect the wound edges together.^86,87^ Therefore, 0° or 180° indicates a fiber bridging the wound and 90° corresponds to a fiber that is perpendicular to wound closure. Initially, fibroblasts on stiffer substrates mimicking fibrotic wounds rapidly assembled a densely aligned EDA Fn matrix (Fig. 2C,D,E) with fiber angles closer to bridging the two edges of the wound (146 kPa: 24h, 112.2°±15.77), than the sparse randomly oriented EDA Fn matrix assembled by fibroblasts on softer substrates mimicking normal wounds (18 kPa: 24h, 93.73°±9.994). Afterwards, fiber angles of EDA Fn matrices hovered around 90°, suggesting that cells switched from bridging the two sides of the wound to filling in the wound with more matrix fibers by assembling matrix fibers laterally (18 kPa: 48h, 85.12°±14.57; 72h, 85.12°±14.57; 146 kPa: 48h, 94.12°±21.91; 72h, 94.12°±21.91).

Collectively, these matrix microarchitecture measurements suggest that fibroblasts on stiffer PDMS substrates, when migrating to close wounds, rapidly assemble a densely aligned EDA Fn matrix network comprising thinner fibers under high tension. The increased mechanical tension in EDA Fn fibers likely contributes to changes in protein conformation or changes to the binding accessibility of the EDA domain for cells.^88–90^ Fibronectin is the first matrix protein deposited by fibroblasts at the wound site, and thus is the predominant matrix protein handling mechanical load during the early wound before collagen replaces the provisional component of the Fn matrix.^27,91^ It is likely EDA Fn possesses a higher mechanical load capacity than EDA^-^ Fn isoforms; the extra

FnIII module, a cryptic binding domain^41^, potentially providing more conformational flexibility to allow fibers to deform or stretch further before resisting cell contractility or microenvironmental forces. Therefore, mechanical modulation can dictate initial EDA Fn matrix microarchitecture within wounds, most prominently in the absence of other matrix elements. Having established that EDA Fn is a key matrix protein in the early stages of our wound assay and its microarchitecture sensitive to mechanical stimuli, we next sought to investigate if inhibiting cell binding to the EDA domain of Fn interferes with overall matrix assembly and microarchitecture during normal and fibrotic wound healing.

### Interfering with cell binding to the EDA domain within Fn

Cell binding to the EDA domain of Fn is responsible for many cellular functions essential to both normal wound healing and fibrotic wound healing.^21,47^ However, how cell adhesion to the EDA domain of Fn contributes to EDA Fn matrix structure-function during wound repair is not fully clear.

Irigenin, a competitive inhibitor of cell binding to the EDA domain within Fn, was exogenously added to the wound assays (Fig. 3A,C).^71^ As Irigenin was reconstituted in dimethylsulfoxide (DMSO), DMSO was added exogenously to parallel wound assays as a control. Moreover, there is a need to quantify early matrix microarchitecture as changes to initial cell binding to EDA Fn and EDA Fn matrix assembly likely dictate the remainder of the wound closure process. Therefore, the remainder of the study focuses on only the 4h and 24h time points.

**Figure 3:**
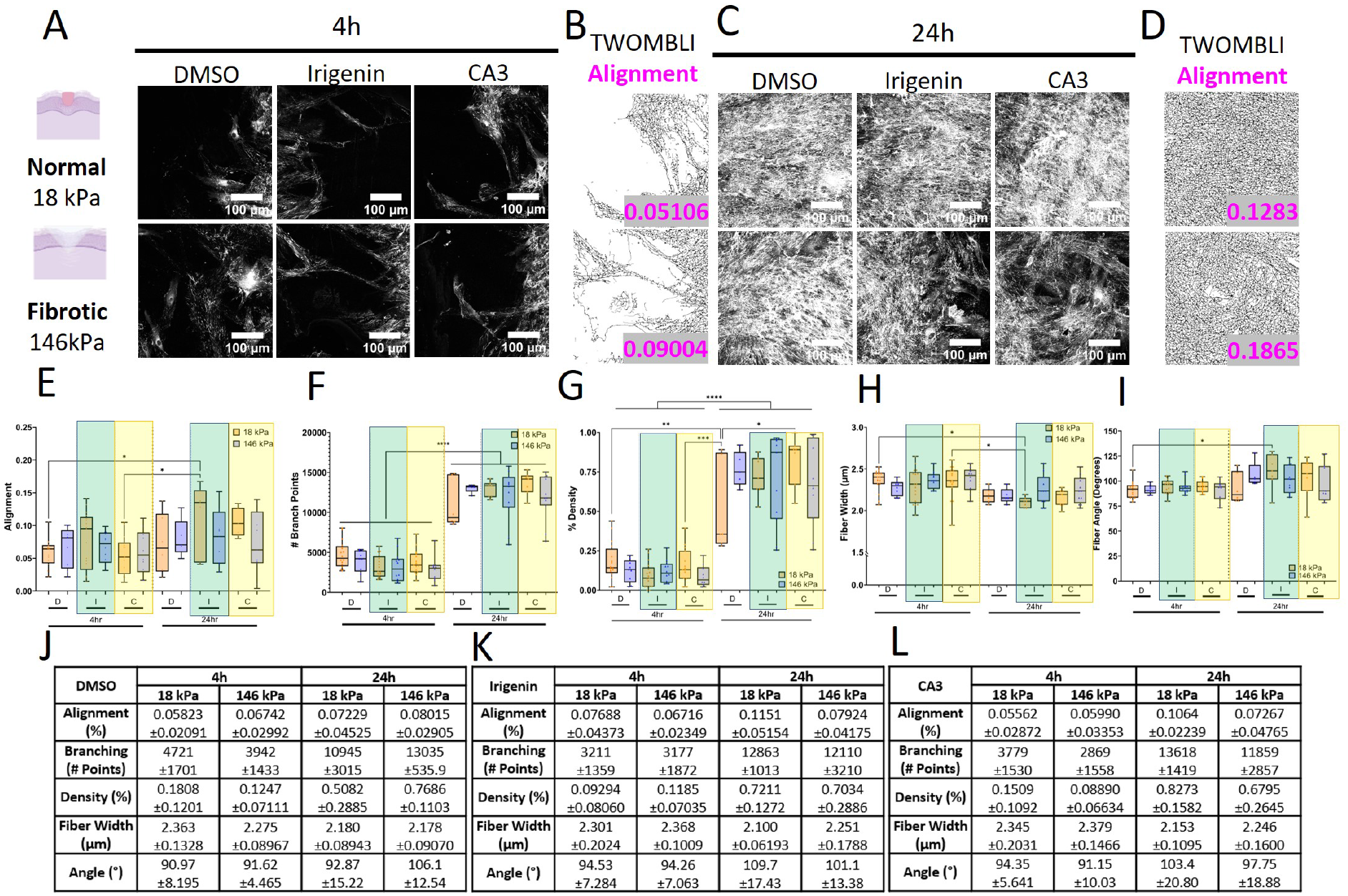
Inhibition of wound EDA fibronectin-mediated matrix-cell binding and partial inhibition of YAP alter EDA Fn matrix microarchitecture over time and stiffness. Fluorescently labeled EDA Fn images of the of the wound assay were taken by confocal microscopy post-wounding at 4h (**A**) and 24h (**C**) of the DMSO control, Irigenin-based inhibition of the EDA Fn mediated matrix-cell binding, and CA3-mediated partial inhibition of YAP. Scale bars 100 µm. **B,D**. Example matrix masks and alignment values generated from our samples by the ImageJ macro, TWOMBLI. **E**. TWOMBLI generated matrix alignment values. *P≤0.0288. **F**. Number of branch points from TWOMBLI analysis of EDA Fn matrices. ****P<0.0001, only same stiffnesses compared across conditions. **G**. EDA Fn matrix density as returned by TWOMBLI analysis. *P=0.0216, **P=0.002, ***P=0.0005, ****P<0.0001, only same stiffnesses selected for comparison across conditions. **H**. Fiber width as returned by CT-FIRE analysis. *P≤0.0282. **I**. Fiber angle from horizontal, CT-FIRE, *P=0.0261. All N≥5, 4h and 24h conditions only, one-way ANOVA, Tukey’s post-hoc analysis. **J,K,L,** Table of mean ± standard deviation for the three test conditions, DMSO (**J**), Irigenin (**K**), and CA3 (**L**), for alignment, branching, density, fiber width, and fiber angle.

First, initial EDA Fn matrices assembled by fibroblasts treated with DMSO (control) migrating into wounds (Fig. 3) reveal similar matrix architectural changes across time and substrate stiffnesses (Fig. 2). Fibroblasts treated with DMSO (control) on stiffer substrates mimicking fibrotic wounds, assembled EDA Fn matrices that were more aligned (146 kPa: 4h, 0.06742±0.02992; 24h, 0.08015±0.02905) than EDA Fn matrices assembled by fibroblasts treated with DMSO (control) on softer substrates mimicking normal wounds (18 kPa: 4h, 0.05823±0.02091; 24h, 0.07229±0.04525) (Fig. 3E,J). However, a new EDA Fn matrix alignment trend emerged when competitively blocking cell binding to the EDA domain of Fn with Irigenin treatment. Fibroblasts treated with Irigenin on stiffer substrates mimicking fibrotic wounds, assembled EDA Fn matrices that were more randomly oriented or less aligned (146 kPa: 4h, 0.06716±0.02349; 24h, 0.07924±0.04175) than EDA Fn matrices assembled by fibroblasts treated with Irigenin on softer substrates mimicking normal wounds (18 kPa: 4h 0.07688±0.04373; 24h, 0.1151±0.05154). This suggests that within fibrotic wounds, treatment with Irigenin, or inhibiting cell binding to the EDA domain of Fn, may restore normal EDA Fn fiber orientation to mitigate the pro-fibrotic response during wound closure. However, within normal wounds, treatment with Irigenin and preventing cell binding to the EDA domain of Fn may unintentionally enhance EDA Fn matrix alignment and drive a pro-fibrotic response; demonstrating that cell binding to the EDA domain of Fn is necessary during normal wound closure.

Then, we examined EDA Fn branching (Fig. 3F,J,K). Fibroblasts treated with DMSO (control) on stiffer substrates mimicking fibrotic wounds, assembled EDA Fn matrices that were initially less branched but then more branched (146 kPa: 4h, 3942±1433; 24h, 13035±535.9) than EDA Fn matrices assembled by fibroblasts treated with DMSO (control) on softer substrates mimicking normal wounds (18 kPa: 4h, 4721±1701; 24h, 10945±3015). Fibroblasts treated with Irigenin on stiffer substrates mimicking fibrotic wounds, assembled EDA Fn matrices with similar levels of matrix branching (146 kPa: 4h, 3177±1872; 24h: 12110±3210) compared with EDA Fn matrices assembled by fibroblasts treated with Irigenin on softer substrates mimicking normal wounds (18 kPa: 4h, 3211±1359; 24h, 12863±1013). Quantifying matrix density (Fig. 3G,J,K) demonstrates that fibroblasts treated with DMSO (control) migrating on stiff substrates mimicking fibrotic wounds (146 kPa: 4h, 0.1247±0.07111; 24h, 0.7686±0.1103) initially assembled less, but then more EDA Fn than that of fibroblasts treated with DMSO (control) migrating on soft substrates mimicking normal wounds (18 kPa: 4h, 0.1808±0.1201; 24h, 0.5082±0.2885). Fibroblasts treated with Irigenin on stiffer substrates mimicking fibrotic wounds, overall assembled similar levels of EDA Fn (146 kPa: 4h, 0.1185 ±0.07035; 24h, 0.7034±0.2886) compared with EDA Fn matrices assembled by fibroblasts treated with Irigenin on softer substrates mimicking normal wounds (18 kPa: 4h, 0.09294±0.08060; 24h, 0.7211±0.1272). As expected, these data indicate that the amount of EDA Fn deposited within normal or fibrotic wounds is unchanged with Irigenin treatment, which would only prevent cell binding to the EDA domain within Fn fibers already assembled.

Next, we quantified EDA Fn fiber width (Fig. 3H,J,K). EDA Fn fibers assembled by fibroblasts treated with DMSO (control) migrating into wounds on stiffer PDMS substrates mimicking fibrotic wounds (146 kPa: 4h, 2.275±0.08967; 24h, 2.178±0.09070) were again thinner than that of EDA Fn fibers assembled by fibroblasts treated with DMSO (control) migrating into wounds on softer PDMS substrates mimicking normal wounds (18 kPA: 4h, 2.363±0.1328; 24h, 2.180±0.08943).

Fibroblasts treated with Irigenin on stiffer substrates mimicking fibrotic wounds, assembled EDA Fn matrices with thicker fibers than that of EDA Fn matrices assembled by fibroblasts treated with Irigenin on softer substrates mimicking normal wounds (18 kPa: 4h, 2.301±0.2024; 24h, 2.100±0.06193. 146 kPa: 4h, 2.368±0.1009; 24h, 2.251±0.1788). We also quantified EDA Fn fiber angles (Fig. 3I,J,K) against the horizontal plane (Fig. 2H). Initially, fibroblasts treated with DMSO (control) on stiffer substrates mimicking fibrotic wounds assembled an aligned EDA Fn matrix (Fig. 3E) with fiber angles around 90° then afterwards fiber angles moving towards 180° (146 kPa: 4h, 91.62°±4.465°; 24h, 106.1°±12.54°), suggesting that cells initially assembled EDA Fn to fill in the wound before switching to bridge the two sides of the fibrotic wound. Fibroblasts treated with DMSO (control) on softer substrates mimicking normal wounds (18 kPa: 4h, 90.97°±8.195°; 24h, 92.87°±15.22°) remained around 90° suggesting that cells were persistently assembling randomly organized matrix fibers laterally to fill in the normal wound. Fibroblasts treated with Irigenin on stiffer substrates mimicking fibrotic wounds assembled a randomly organized EDA Fn matrix (Fig. 3E) with fiber angles persistently around 90°, suggesting that inhibiting cell binding to the EDA domain of Fn led cells to continue filling in the fibrotic Irigenin-treated wound. Fibroblasts treated with Irigenin on softer substrates mimicking normal wounds, however, assembled a more aligned EDA Fn matrix (Fig. 3D) with fiber angles initially around 90° then moving towards 180°, suggesting that cells initially assembled EDA Fn to fill in the wound before switching to bridge the two sides of the normal wound. This suggests that within fibrotic wounds, treatment with Irigenin, likely provides a stress-shielding effect as prevent cells from binding to the EDA domain of Fn may restore normal EDA Fn fiber tension and organization to mitigate the pro-fibrotic response during wound closure. However, within normal wounds, treatment with Irigenin may unintentionally enhance EDA Fn fiber tension and dysregulate matrix organization to drive a pro-fibrotic response.

Collectively, these matrix microarchitecture measurements suggest that fibroblasts on stiffer PDMS substrates, when treated with Irigenin to prevent cell binding to the EDA domain within Fn, might restore normal wound closure processes by assembling a randomly oriented EDA Fn matrix network comprising thicker fibers likely under less tension. The EDA domain within Fn may play a role in mediating the degree to which cells sense matrix tension; reducing it during normal wound healing but enhancing it during fibrotic wound healing.

### Potential cross-talk between EDA Fn organization and YAP activity

Subsequently, to evaluate the underlying mechanism by which fibroblasts are responding and maintaining the different organization of EDA Fn within normal and fibrotic wounds, we looked to YAP as a potential downstream effector of EDA Fn (Fig. 3L). YAP is a mechanotransduction signaling effector that not only is essential for wound closure^53^ and associated with tissue regeneration,^45,58,59^ but also is directly correlated with tissue fibrosis.^92^ EDA Fn shares these same contradictory effects^21,43–50^ on wound healing.^20,21^ Given that YAP can be activated by not only substrate stiffness^93^ and geometry^94,95^ but also by ECM proteins including Fn,^53,56^ YAP became a leading candidate for investigation with respect to EDA Fn matrix organization in wound healing. We exogenously delivered CA3 to our wound assays to attenuate YAP activity. Complete inhibition of YAP was not desired as YAP is essential for wound closure; therefore a non-toxic (0.5 µM/mL) concentration of CA3 was used.^62^

Fibroblasts treated with CA3 on stiffer substrates mimicking fibrotic wounds, though initially did not appear to fully alter EDA Fn matrix assembly and microarchitecture at 4h, did assemble EDA Fn matrices that by 24h, were overall less aligned, less branched, less dense, and comprised thicker fibers primarily oriented laterally to the wound edge than EDA Fn matrices assembled by fibroblasts treated with CA3 on softer substrates (Fig. 3L). These data suggest that mitigating YAP signaling in cells on stiffer substrates mimicking fibrotic wounds might require time for cells to respond and restore normal matrix assembly and wound closure mechanisms. Given that fibroblasts treated with Irigenin or CA3 on fibrotic substrates similarly assemble normal EDA Fn matrix microarchitecture by 24h, there is likely a mechanoregulatory link between EDA Fn structure-function and YAP activity that drives normal wound healing. Mechanistically, YAP activation requires cytoplasmic YAP to translocate into the nucleus to alter target gene expression.^63^ Therefore, we next measured the nuclear/cytoplasmic ratio of YAP to investigate YAP activity within fibroblasts migrating on PDMS substrates mimicking normal and fibrotic wounds.

### YAP differentially activated by substrate stiffness

We evaluated initial YAP activity of fibroblasts just beginning to migrate into wounds (4h) and later YAP activity of fibroblasts fully migrated into wounds (24h) (Fig. 4). Fibroblasts treated with DMSO (control) on stiffer substrates mimicking fibrotic wounds, decreased YAP activation (Fig. 4C,E: 146 kPa: 4h, 1.631±0.3558; 24h, 1.54±0.4746), compared to fibroblasts treated with DMSO (control) on softer substrates mimicking normal wounds (18 kPa: 4h, 1.939±0.4037, 24h, 1.74±0.3906). YAP is required for many functions involved in wound healing, including migration and proliferation. A recent study suggests that YAP activity is dependent on adhesion maturation and a balance of forces between cell contractility and matrix rigidity.^96^ Given that EDA Fn can likely bear more mechanical load and when assembled within normal wounds with random orientation and thicker fibers, it seems likely that fibroblasts are able to slowly reinforce and mature focal adhesions, potentially modifying their mechanosensing to translocate YAP into the nucleus. Additionally, focal adhesion maturation can be dependent on surface topography or ligand density,^97–101^ which in our wound assay system would suggest focal adhesion maturation could be dependent on fiber width. In our fibrotic wounds, EDA Fn is more aligned and comprises thinner fibers, which likely limits the size of focal adhesions and potentially limits YAP activity. Therefore, it is unsurprising that fibroblasts nuclear translocates more YAP in normal wounds.

**Figure 4:**
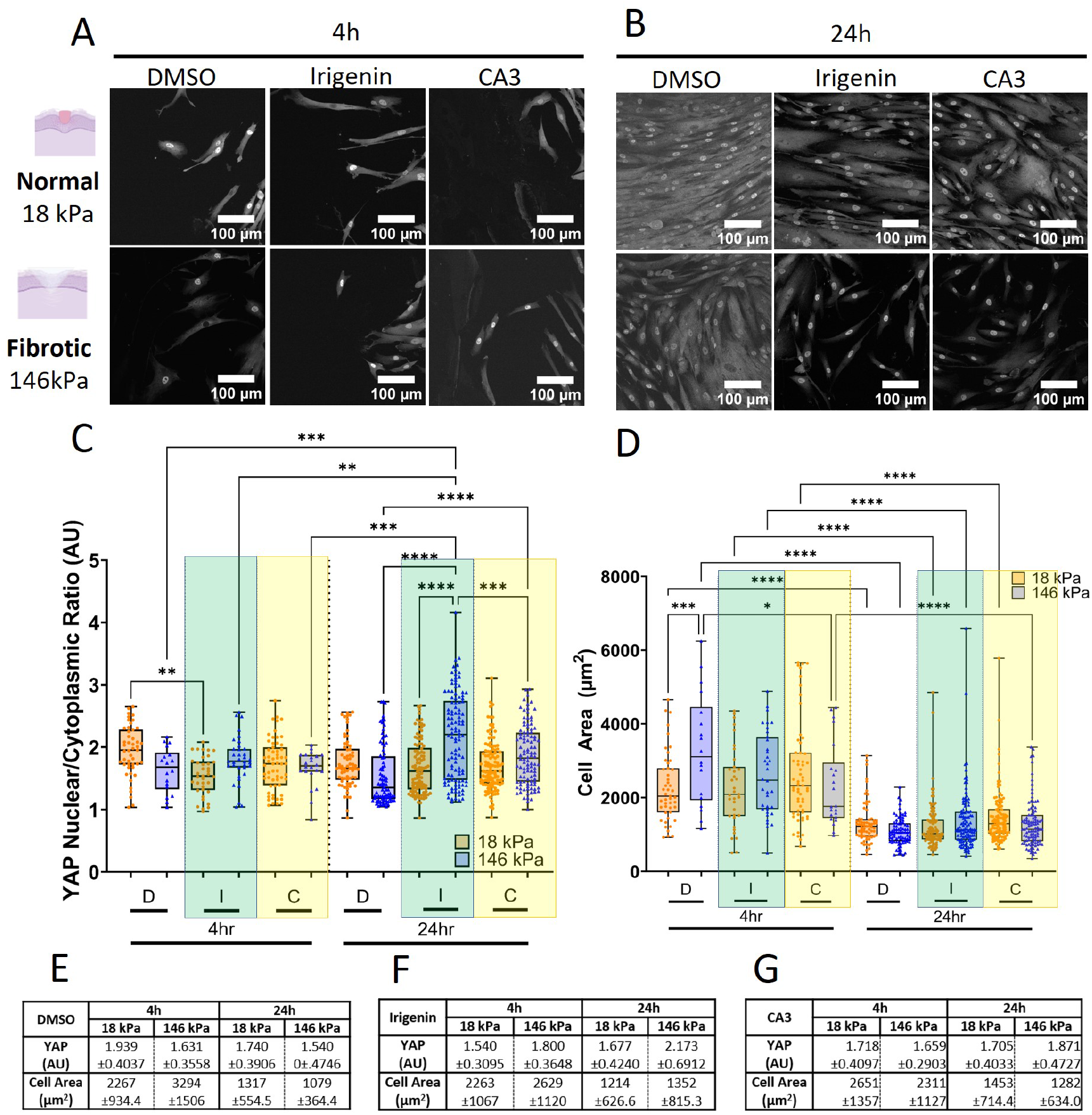
Stiffness alters 4h Irigenin and CA3 YAP activation and cell area; the 24h YAP profile affected by contact inhibition. **A,B.** Representative 4h and 24h fluorescently labeled YAP images were taken by confocal microscopy and analyzed prior to image adjustment for presentation (brightness +40% and contrast -40%). Scale bars 100 µm. **C**. 4h and 24h nuclear/cytoplasmic YAP analysis, selected significances: *P=0.0153, **P=0.0057, ***P≤0.003, ****P<0.0001. **D**. 4h and 24h cell area analysis. *P=0.0103 , ***P=0.005, ****P<0.0001, horizontal line shows comparison of same stiffnesses across time points at ****P<0.0001. All N≥18, one-way ANOVA, Tukey’s post-hoc analysis. **E,F,G**. Table of mean ± standard deviation for the three test conditions, DMSO (**E**), Irigenin (**F**), and CA3 (**G**), for YAP nuclear/cytoplasmic ratio and cell area.

Interestingly, when fibroblasts on stiffer substrates were treated with Irigenin to inhibit cell binding to the EDA domain within Fn, their YAP activity increased (146 kPa: 4h, 1.8±0.3648; 24h, 2.173±0.6912) compared to the YAP activity of fibroblasts treated with DMSO (control) on the same stiffness (146 kPa). Similar increases to YAP activity were quantified even when fibroblasts on stiffer substrates were treated with CA3 to mitigate YAP activity (146 kPa: 4h, 1.659±0.2903; 24h, 1.871±0.4727) compared to the YAP activity of fibroblasts treated with DMSO (control) on the same stiffness (146 kPa). It is likely that changes to EDA Fn microarchitecture, when assembled within fibrotic wounds and treated with Irigenin or CA3, now more randomly organized comprising thicker fibers, allow focal adhesion maturation and therefore increases YAP activity. When inhibiting cell binding to the EDA domain within Fn or when mitigating YAP activity by fibroblasts on soft substrates mimicking normal wounds, there was similarly reduced nuclear YAP translocation (Irigenin: 4h, 1.54±0.3095; 24h, 1.677±0.4240; CA3: 4h, 1.718±0.4097; 24h, 1.705±0.0.4033) compared to nuclear YAP translocation by fibroblasts treated with DMSO (control) on the same stiffness (18 kPa). Suggesting that cell binding to the EDA domain within Fn regulates YAP activation in normal wounds and that a certain level of YAP activity is necessary for normal EDA Fn assembly and organization. Taken together, these results indicate a potential mechanism to mitigate stiffness-mediated pro-fibrotic responses.

Interestingly and contrary to our results, increased fibrotic tissue stiffness has typically been shown to drive pathological YAP activation.^6^ However, depending on various factors including cell shape, topography,^95^ and substrate viscoelasticity,^102^ force is not always directly correlated to increased YAP activation.^103^ Viscoelasticity alters the mechanical regulation of cells.^68^ Closer examination of viscoelastic contributions to YAP signaling reveals a complex YAP activation profile deeply impacted by local microenvironmental forces. Viscoelastic substrates encourage YAP translocation into the nucleus compared to elastic substrates.^70^ Further, fast-relaxing viscoelastic material encourages more YAP translocation into the nucleus than slow-relaxing viscoelastic substrates.^70,104^ Similarly supported by our YAP results when fibroblasts are treated with DMSO as a control, others show that treatment with DMSO leads to YAP activating in an inverse stiffness-dependent manner on a substrate stiffness ranging from 5 kPa to 75 kPa.^105^ Conversely, from 1 kPa to 40 kPa, it has been previously demonstrated that increased nuclear localization of YAP is directly correlated with increasing stiffness.^56^ In our wound assays, when fibroblasts were treated with Irigenin or CA3, nuclear localization of YAP is directly correlated with substrate stiffness. The discrepancy within literature and our results suggest that YAP activation is likely further regulated by matrix structure. In normal wounds when EDA Fn more randomly oriented with thicker fibers and likely under less tension allowing for focal adhesion maturation, YAP activity is high. But in fibrotic wounds when EDA Fn is highly aligned with thinner fibers and likely under more tension focal adhesion maturation processes may be disrupted, YAP activity is lower. The complex, nuanced responses of YAP to various mechanical cues demonstrates the need to better understand and contextualize matrix mediated YAP signaling.

In addition to migration, YAP independently regulates cell size ^106–108^ as cell area is directly associated with cell attachment.^109^ Therefore, we next quantified fibroblast cell area to further contextualize our nuclear YAP localization results (Fig. 4D). Initially, fibroblasts treated with DMSO (control) on stiffer substrates mimicking fibrotic wounds were more spread (4h, 3294μm^2^±1506) than fibroblasts treated with DMSO (control) on softer substrates mimicking normal wounds (4h, 2267μm^2^±934.4). However, by 24h when the wound was mostly closed, fibroblasts treated with DMSO (control) on stiffer substrates mimicking fibrotic wounds were less spread (24h, 1079μm^2^±364.4) than fibroblasts treated with DMSO (control) on softer substrates mimicking normal wounds (4h, 1317μm^2^±554.5). This suggests that fibroblasts on stiffer substrates mimicking fibrotic wounds are quicker to alter their behavior from being more spread and proliferative (filling in wound with granulation tissue) and switching^110^ to enhancing matrix deposition (fibrosis) than fibroblasts on softer substrates mimicking normal wounds. Additionally, both Irigenin and CA3 treated fibroblasts on stiffer substrates mimicking fibrotic wounds were initially less spread (Irigenin: 4h, 2629μm^2^±1120; CA3: 4h, 2311μm^2^±1127) than fibroblasts treated with DMSO (control) on the same stiffness (146 kPa, 4h). However, afterwards, both Irigenin and CA3 treated fibroblasts on stiffer substrates mimicking fibrotic wounds were more spread (Irigenin: 24h, 1352μm^2^±815.3; CA3: 24h, 1282μm^2^±634) than fibroblasts treated with DMSO (control) on the same stiffness (146 kPa, 24h). Taken together with the increase in YAP activity, it is likely that these Irigenin and CA3 treated fibroblasts are likely proliferating to fill in a wound with granulation tissue. This suggests that changes to EDA Fn microarchitecture now more randomly organized comprising thicker fibers under less tension, potentially decreases the rigidity sensed by fibroblasts even on stiffer substrates mimicking fibrotic wounds leading to smaller cells;^111^ further highlighting Irigenin and CA3 as potential tools for mitigating the initial pro-fibrotic cascade.

## Conclusion

Here, we developed a 2D wound closure assay to investigate the effects of substrate stiffness mimicking normal or fibrotic wounds on fibroblast assembled EDA Fn microarchitecture and YAP signaling during wound healing. On soft substrates mimicking normal wounds, EDA Fn was randomly organized comprising thicker fibers which led to increased YAP activity. On stiffer substrates mimicking fibrotic wounds, EDA Fn was highly aligned comprising thinner fibers which likely disrupted YAP activation. Our wound closure assays also demonstrated that preventing cell binding to the EDA domain within Fn fibers or attenuating YAP activity both similarly led to less aligned EDA Fn comprising thicker fibers and increased YAP activity (Fig. 5). These results demonstrate the potential in controlling initial EDA Fn matrix organization and regulating YAP activity to ensure normal wound repair processes. Our wound closure assay will enable future studies to investigate how EDA Fn fiber structure might alter cell adhesion and migration, and furthermore, inform therapeutic strategies to control wound repair mechanisms.

**Figure 5:**
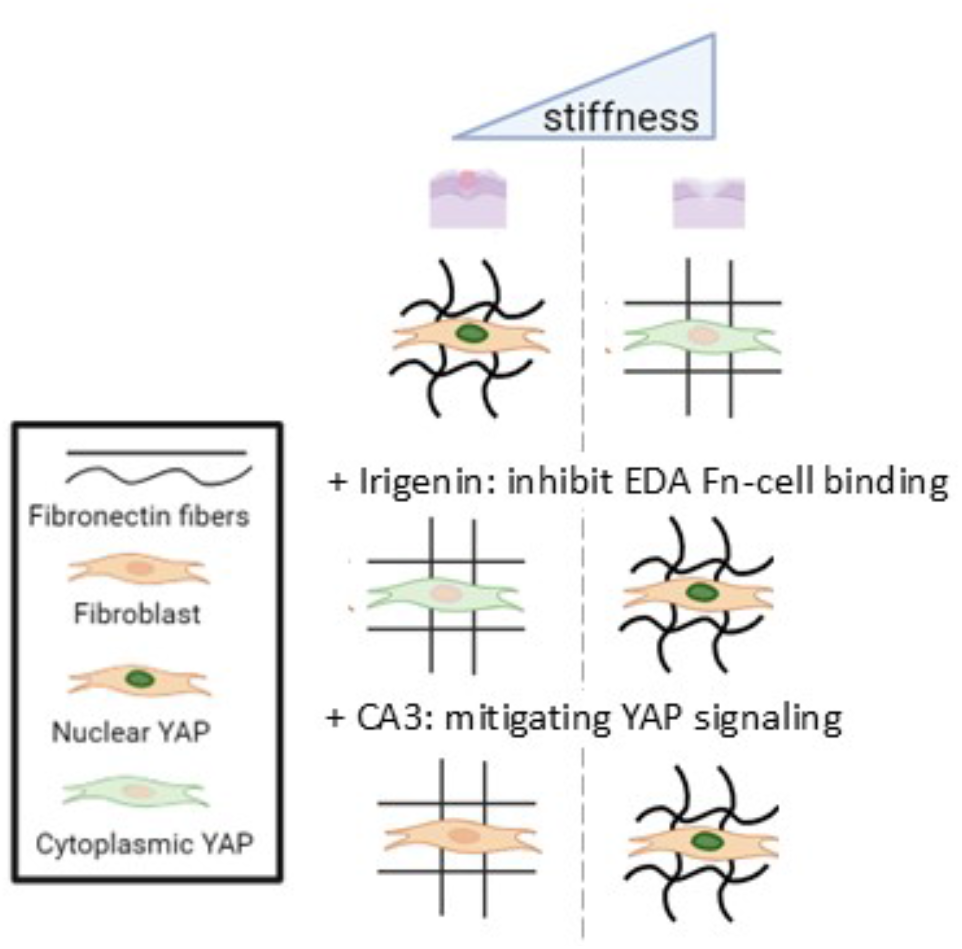
Schematic illustrating changes in cell and matrix responses during normal and fibrotic wound closure. Left column: soft substrates mimicking normal wounds (18 kPa). Right column: stiff substrates mimicking fibrotic wounds (146 kPa). Representative schematic of EDA Fn fiber width, alignment, and YAP localization for control (1^st^ row), Irigenin-treated wound closure (2^nd^ row), and CA3-treated wound closure (3^rd^ row).

## Acknowledgments

K.W. acknowledges startup funds from the Bioengineering Department at Temple University and a grant (#2305) from the W.W. Smith Charitable Trust. J.D.B was supported by the National Institute of Health (5T34 GM 136494). The authors thank the shared departmental resources at Temple University for equipment usage. We thank BioRender for providing a platform to create the schematics used in figures.

## Credit Author Statement

**Jennifer Patten**: Conceptualization, Methodology, Validation, Data Analysis, Visualization, Writing. **Patrick Halligan**, Methodology, Validation, Data Analysis, Writing. **Ghazal Bashiri**: Review and Editing. **Michael Kegel**: Methodology, Validation. **Jacob Bonadio**: Methodology, Review. **Karin Wang**: Conceptualization, Methodology, Validation, Visualization, Writing, Funding acquisition, Supervision.

## References

1 Verhaegen PDHM, van Zuijlen PPM, Pennings NM, van Marle J, Niessen FB, van der Horst CMAM, et al. Differences in collagen architecture between keloid, hypertrophic scar, normotrophic scar, and normal skin: An objective histopathological analysis. Wound Repair Regen 2009;17:649–56. 10.1111/j.1524-475X.2009.00533.x.

2 Garrison CM, Schwarzbauer JE. Fibronectin fibril alignment is established upon initiation of extracellular matrix assembly. Mol Biol Cell 2021;32:739–52. 10.1091/mbc.e20-08-0533.

3 Patten J, Wang K. Fibronectin in development and wound healing. Adv Drug Deliv Rev 2021;170:353–68. 10.1016/j.addr.2020.09.005.

4 Li B, Moshfegh C, Lin Z, Albuschies J, Vogel V. Mesenchymal stem cells exploit extracellular matrix as mechanotransducer. Sci Rep 2013;3:1–8. 10.1038/srep02425.

5 Wang WY, Pearson AT, Kutys ML, Choi CK, Wozniak MA, Baker BM, et al. Extracellular matrix alignment dictates the organization of focal adhesions and directs uniaxial cell migration. APL Bioeng 2018;2:. 10.1063/1.5052239.

6 Noguchi S, Saito A, Nagase T. YAP/TAZ signaling as a molecular link between fibrosis and cancer. Int J Mol Sci 2018;19:. 10.3390/ijms19113674.

7 Kenny FN, Drymoussi Z, Delaine-Smith R, Kao AP, Laly AC, Knight MM, et al. Tissue stiffening promotes keratinocyte proliferation through activation of epidermal growth factor signaling. J Cell Sci 2018;131:. 10.1242/jcs.215780.

8 Eckes B, Nischt R, Krieg T. Cell-matrix interactions in dermal repair and scarring. Fibrogenes Tissue Repair 2010;3:1–11. 10.1186/1755-1536-3-4.

9 Naba A. Mechanisms of assembly and remodelling of the extracellular matrix. Nat Rev Mol Cell Biol 2024. 10.1038/s41580-024-00767-3.

10 Henderson NC, Rieder F, Wynn TA. Fibrosis: from mechanisms to medicines. Nature 2020;587:555–66. 10.1038/s41586-020-2938-9.

11 Dohi T, Padmanabhan J, Akaishi S, Than PA, Terashima M, Matsumoto NN, et al. The Interplay of Mechanical Stress, Strain, and Stiffness at the Keloid Periphery Correlates with Increased Caveolin-1/ROCK Signaling and Scar Progression. Plast Reconstr Surg 2019;144:58e–67e. 10.1097/PRS.0000000000005717.

12 Ito M, Yang Z, Andl T, Cui C, Kim N, Millar SE, et al. Wnt-dependent de novo hair follicle regeneration in adult mouse skin after wounding. Nature 2007;447:316–20. 10.1038/nature05766.

13 Chen K, Kwon SH, Henn D, Kuehlmann BA, Tevlin R, Bonham CA, et al. Disrupting biological sensors of force promotes tissue regeneration in large organisms. Nat Commun 2021;12:1–15. 10.1038/s41467-021-25410-z.

14 Chen K, Henn D, Januszyk M, Barrera JA, Noishiki C, Bonham CA, et al. Disrupting mechanotransduction decreases fibrosis and contracture in split-thickness skin grafting. Sci Transl Med 2022;14:1–18. 10.1126/scitranslmed.abj9152.

15 Leahy DJ, Aukhil I, Erickson HP. 2.0 Å crystal structure of a four-domain segment of human fibronectin encompassing the RGD loop and synergy region. Cell 1996;84:155–64. 10.1016/S0092-8674(00)81002-8.

16 Potts JR. Structure and Assembly. Struct Assem 1975:648–55. 10.1007/978-1-4684-2709-7.

17 Rounsevell RWS, Clarke J. FnIII Domains: Predicting Mechanical Stability. Structure 2004;12:4–5. 10.1016/j.str.2003.12.006.

18 Krammer A, Lu H, Isralewitz B, Schulten K, Vogel V. Forced unfolding of the fibronectin type III module reveals a tensile molecular recognition switch. Proc Natl Acad Sci U S A 1999;96:1351–6. 10.1073/pnas.96.4.1351.

19 Gao M, Craig D, Krammer A, Puklin-Faucher E, Lu H, Vogel V, et al. Fibronectin and Integrin. Theoretical and Computational Biophysics Group. 2006. URL: https://www.ks.uiuc.edu/Research/fibronectin/.

20 To WS, Midwood KS. Plasma and cellular fibronectin: Distinct and independent functions during tissue repair. Fibrogenes Tissue Repair 2011;4:21. 10.1186/1755-1536-4-21.

21 Lenselink EA. Role of fibronectin in normal wound healing. Int Wound J 2015;12:313–6. 10.1111/iwj.12109.

22 Jarnagin WR, Rockey DC, Koteliansky VE, Wang S, Bissell DM. Expression of Variant Fibronectins in Wound Healing: Cellular Source and Biological Activity of the EIIIA Segment in Rat Hepatic Fibrogenesis 2013;127:1–12.

23 Gui L, Wojciechowski K, Gildner CD, Nedelkovska H, Hocking DC. Identification of the heparin-binding determinants within fibronectin repeat III1: Role in cell spreading and growth. J Biol Chem 2006;281:34816–25. 10.1074/jbc.M608611200.

24 Lickert S, Kenny M, Selcuk K, Mehl JL, Bender M, Früh SM, et al. Platelets drive fibronectin fibrillogenesis using integrin αIIbβ3. Sci Adv 2022;8:1–17. 10.1126/sciadv.abj8331.

25 Chen LB, Murray A, Segal RA, Bushnell A, Walsh ML. Studies on intercellular LETS glycoprotein matrices. Cell 1978;14:377–91. 10.1016/0092-8674(78)90123-X.

26 Singer II. The fibronexus: a transmembrane association of fibronectin-containing fibers and bundles of 5 nm microfilaments in hamster and human fibroblasts. Cell 1979;16:675–85. 10.1016/0092-8674(79)90040-0.

27 Wang K, Andresen Eguiluz RC, Wu F, Seo BR, Fischbach C, Gourdon D. Stiffening and unfolding of early deposited-fibronectin increase proangiogenic factor secretion by breast cancer-associated stromal cells. Biomaterials 2015;54:63–71. 10.1016/j.biomaterials.2015.03.019.

28 Sun BK, Siprashvili Z, Khavari PA. Advances in skin grafting and treatment of cutaneous wounds. Science (80- ) 2014;346:941–5. 10.1126/science.1253836.

29 Singh P, Carraher C, Schwarzbauer JE. Assembly of Fibronectin Extracellular Matrix. Annu Rev Cell Dev Biol 2010;26:397–419. 10.1146/annurev-cellbio-100109-104020.

30 Carraher CL, Schwarzbauer JE. Regulation of matrix assembly through rigidity-dependent fibronectin conformational changes. J Biol Chem 2013;288:14805–14. 10.1074/jbc.M112.435271.

31 Kliewe F, Kaling S, Lötzsch H, Artelt N, Schindler M, Rogge H, et al. Fibronectin is up-regulated in podocytes by mechanical stress. FASEB J 2019:fj.201900978RR. 10.1096/fj.201900978RR.

32 Yeh YC, Corbin EA, Caliari SR, Ouyang L, Vega SL, Truitt R, et al. Mechanically dynamic PDMS substrates to investigate changing cell environments. Biomaterials 2017;145:23–32. 10.1016/j.biomaterials.2017.08.033.

33 Efthymiou G, Radwanska A, Grapa AI, Beghelli-De la Forest Divonne S, Grall D, Schaub S, et al. Fibronectin Extra Domains tune cellular responses and confer topographically distinct features to fibril networks. J Cell Sci 2021;134:. 10.1242/jcs.252957.

34 Goossens K, Van Soom A, Van Zeveren A, Favoreel H, Peelman LJ. Quantification of Fibronectin 1 (FN1) splice variants, including two novel ones, and analysis of integrins as candidate FN1 receptors in bovine preimplantation embryos. BMC Dev Biol 2009;9:1–16. 10.1186/1471-213X-9-1.

35 Tomasek JJ, Gabbiani G, Hinz B, Chaponnier C, Brown RA. Myofibroblasts and mechano: Regulation of connective tissue remodelling. Nat Rev Mol Cell Biol 2002;3:349–63. 10.1038/nrm809.

36 Shinde A V., Bystroff C, Wang C, Vogelezang MG, Vincent PA, Hynes RO, et al. Identification of the peptide sequences within the EIIIA (EDA) segment of fibronectin that mediate integrin α9β1- dependent cellular activities. J Biol Chem 2008;283:2858–70. 10.1074/jbc.M708306200.

37 Rossnagl S, Altrock E, Sens C, Kraft S, Rau K, Milsom MD, et al. EDA-Fibronectin Originating from Osteoblasts Inhibits the Immune Response against Cancer. PLoS Biol 2016;14:1–32. 10.1371/journal.pbio.1002562.

38 Arslan F, Smeets MB, Riem Vis PW, Karper JC, Quax PH, Bongartz LG, et al. Lack of fibronectin-EDA promotes survival and prevents adverse remodeling and heart function deterioration after myocardial infarction. Circ Res 2011;108:582–92. 10.1161/CIRCRESAHA.110.224428.

39 Kelsh-Lasher RM, Ambesi A, Bertram C, McKeown-Longo PJ. Integrin α4β1 and TLR4 Cooperate to Induce Fibrotic Gene Expression in Response to Fibronectin’s EDA Domain. J Invest Dermatol 2017;137:2505–12. 10.1016/j.jid.2017.08.005.

40 Lemanska-Perek A, Adamik B. Fibronectin and its soluble EDA-FN isoform as biomarkers for inflammation and sepsis. Adv Clin Exp Med 2019;28:1561–7. 10.17219/acem/104531.

41 Julier Z, Martino MM, De Titta A, Jeanbart L, Hubbell JA. The TLR4 agonist fibronectin extra domain a is cryptic, Exposed by elastase-2; Use in a fibrin matrix cancer vaccine. Sci Rep 2015;5:1–10. 10.1038/srep08569.

42 Okamura Y, Watari M, Jerud ES, Young DW, Ishizaka ST, Rose J, et al. The Extra Domain A of Fibronectin Activates Toll-like Receptor 4. J Biol Chem 2001;276:10229–33. 10.1074/jbc.M100099200.

43 Muro AF, Chauhan AK, Gajovic S, Iaconcig A, Porro F, Stanta G, et al. Regulated splicing of the fibronectin EDA exon is essential for proper skin wound healing and normal lifespan. J Cell Biol 2003;162:149–60. 10.1083/jcb.200212079.

44 Klingberg F, Chau G, Walraven M, Boo S, Koehler A, Chow ML, et al. The fibronectin ED-A domain enhances recruitment of latent TGF-β-binding protein-1 to the fibroblast matrix. J Cell Sci 2018;131:1–12. 10.1242/jcs.201293.

45 Brewer CM, Nelson BR, Wakenight P, Collins S, Daryl M, Dong XR, et al. Adaptations in Hippo-Yap signaling and myofibroblast fate underlie scar-free ear appendage wound healing in spiny mice 2021;56:2722–40. 10.1016/j.devcel.2021.09.008.Adaptations.

46 Dobaczewski M, Gonzalez-Quesada C, Frangogiannis NG. The extracellular matrix as a modulator of the inflammatory and reparative response following myocardial infarction. J Mol Cell Cardiol 2010;48:504–11. 10.1016/j.yjmcc.2009.07.015.

47 Bhattacharyya S, Tamaki Z, Wang W, Hinchcliff M, Hoover P, Getsios S, et al. FibronectinEDA promotes chronic cutaneous fibrosis through toll-like receptor signaling. Sci Transl Med 2014;6:. 10.1126/scitranslmed.3008264.

48 Cooper JG, Jeong SJ, McGuire TL, Sharma S, Wang W, Bhattacharyya S, et al. Fibronectin EDA forms the chronic fibrotic scar after contusive spinal cord injury. Neurobiol Dis 2018;116:60–8. 10.1016/j.nbd.2018.04.014.

49 Bhattacharyya S, Wang W, Tamaki Z, Shi B, Yeldandi A, Tsukimi Y, et al. Pharmacological inhibition of tolllike receptor-4 signaling by TAK242 prevents and induces regression of experimental organ fibrosis. Front Immunol 2018;9:1–10. 10.3389/fimmu.2018.02434.

50 Tschumperlin DJ, Lagares D. Mechano-therapeutics: Targeting Mechanical Signaling in Fibrosis and Tumor Stroma. Pharmacol Ther 2020;212:107575. 10.1016/j.pharmthera.2020.107575.

51 Velling T, Risteli J, Wennerberg K, Mosher DF, Johansson S. Polymerization of type I and III collagens is dependent on fibronectin and enhanced by integrins α11β1 and α2β1. J Biol Chem 2002;277:37377–81. 10.1074/jbc.M206286200.

52 Sottile J, Hocking DC. Fibronectin Polymerization Regulates the Composition and Stability of Extracellular Matrix Fibrils and Cell-Matrix Adhesions Jane. Mol Biol Cell 2002;13:3546–59. 10.1091/mbc.E02.

53 Liu F, Lagares D, Choi KM, Stopfer L, Marinković A, Vrbanac V, et al. Mechanosignaling through YAP and TAZ drives fibroblast activation and fibrosis. Am J Physiol - Lung Cell Mol Physiol 2015;308:L344–57. 10.1152/ajplung.00300.2014.

54 Cai X, Wang KC, Meng Z. Mechanoregulation of YAP and TAZ in Cellular Homeostasis and Disease Progression. Front Cell Dev Biol 2021;9:1–12. 10.3389/fcell.2021.673599.

55 Aragona M, Panciera T, Manfrin A, Giulitti S, Michielin F, Elvassore N, et al. A mechanical checkpoint controls multicellular growth through YAP/TAZ regulation by actin-processing factors. Cell 2013;154:1047–59. 10.1016/j.cell.2013.07.042.

56 Dupont S, Morsut L, Aragona M, Enzo E, Giulitti S, Cordenonsi M, et al. Role of YAP/TAZ in mechanotransduction. Nature 2011;474:179–84. 10.1038/nature10137.

57 Van Der Stoel M, Schimmel L, Nawaz K, Van Stalborch AM, De Haan A, Klaus-Bergmann A, et al. DLC1 is a direct target of activated YAP/TAZ that drives collective migration and sprouting angiogenesis. J Cell Sci 2020;133:. 10.1242/jcs.239947.

58 Grzelak EM, Elshan, NGR DayanShao S, Bulos ML, Joseph SB, Chatterjee AK, Chen JJ, et al. Pharmacological YAP activation promotes regenerative repair of cutaneous wounds. Proc Natl Acad Sci 2023;120:. 10.1073/pnas.

59 Moya IM, Halder G. Hippo–YAP/TAZ signalling in organ regeneration and regenerative medicine. Nat Rev Mol Cell Biol 2019;20:211–26. 10.1038/s41580-018-0086-y.

60 Hu M, Wang H, Li S, Yan F, Fu C, Li L, et al. Yes-associated protein is involved in myocardial fibrosis in rats with diabetic cardiomyopathy. Diabetes, Metab Syndr Obes Targets Ther 2021;14:2133–43. 10.2147/DMSO.S302466.

61 Chaudhuri O, Gu L, Darnell M, Klumpers D, Bencherif SA, Weaver JC, et al. Substrate stress relaxation regulates cell spreading. Nat Commun 2015;6:1–7. 10.1038/ncomms7365.

62 Song S, Xie M, Scott AW, Jin J, Ma L, Dong X, et al. A novel YAP1 inhibitor targets CSC-enriched radiation-resistant cells and exerts strong antitumor activity in esophageal adenocarcinoma. Mol Cancer Ther 2018;17:443–54. 10.1158/1535-7163.MCT-17-0560.

63 Piersma B, Bank RA, Boersema M. Signaling in fibrosis: TGF-β, WNT, and YAP/TAZ converge. Front Med 2015;2:1–14. 10.3389/fmed.2015.00059.

64 Kollmannsberger P, Bidan CM, Dunlop JWC, Fratzl P, Vogel V. Tensile forces drive a reversible fibroblast-to-myofibroblast transition during tissue growth in engineered clefts. Sci Adv 2018;4:. 10.1126/sciadv.aao4881.

65 Lamar JM, Xiao Y, Norton E, Jiang ZG, Gerhard GM, Kooner S, et al. SRC tyrosine kinase activates the YAP/TAZ axis and thereby drives tumor growth and metastasis. J Biol Chem 2019;294:2302–17. 10.1074/jbc.RA118.004364.

66 Bugg D, Bretherton R, Kim P, Olszewski E, Nagle A, Schumacher AE, et al. Infarct Collagen Topography Regulates Fibroblast Fate via p38-Yes-Associated Protein Transcriptional Enhanced Associate Domain Signals. Circ Res 2020;127:1306–22. 10.1161/CIRCRESAHA.119.316162.

67 Mascharak S, Talbott HE, Januszyk M, Griffin M, Chen K, Davitt MF, et al. Multi-omic analysis reveals divergent molecular events in scarring and regenerative wound healing. Cell Stem Cell 2022;29:315–327.e6. 10.1016/j.stem.2021.12.011.

68 Dwivedi N, Das S, Bellare J, Majumder A. Viscoelastic substrate decouples cellular traction force from other related phenotypes. Biochem Biophys Res Commun 2021;543:38–44. 10.1016/j.bbrc.2021.01.027.

69 Cameron AR, Frith JE, Gomez GA, Yap AS, Cooper-White JJ. The effect of time-dependent deformation of viscoelastic hydrogels on myogenic induction and Rac1 activity in mesenchymal stem cells. Biomaterials 2014;35:1857–68. 10.1016/j.biomaterials.2013.11.023.

70 Elosegui-Artola A, Gupta A, Najibi AJ, Seo BR, Garry R, Tringides CM, et al. Matrix viscoelasticity controls spatiotemporal tissue organization. Nat Mater 2023;22:117–27. 10.1038/s41563-022-01400-4.

71 Amin A, Chikan NA, Mokhdomi TA, Bukhari S, Koul AM, Shah BA, et al. Irigenin, a novel lead from Western Himalayan chemiome inhibits Fibronectin-Extra Domain A induced metastasis in Lung cancer cells. Sci Rep 2016;6:1–13. 10.1038/srep37151.

72 Wershof E, Park D, Barry DJ, Jenkins RP, Rullan A, Wilkins A, et al. A FIJI macro for quantifying pattern in extracellular matrix. Life Sci Alliance 2021;4:1–11. 10.26508/LSA.202000880.

73 Bredfeldt JS, Liu Y, Pehlke CA, Conklin MW, Szulczewski JM, Inman DR, et al. Computational segmentation of collagen fibers from second-harmonic generation images of breast cancer. J Biomed Opt 2014;19:016007. 10.1117/1.jbo.19.1.016007.

74 Masson-Meyers DS, Andrade TAM, Caetano GF, Guimaraes FR, Leite MN, Leite SN, et al. Experimental models and methods for cutaneous wound healing assessment. Int J Exp Pathol 2020;101:21–37. 10.1111/iep.12346.

75 Jonkman JEN, Cathcart JA, Xu F, Bartolini ME, Amon JE, Stevens KM, et al. Cell Adhesion & Migration An introduction to the wound healing assay using livecell microscopy An introduction to the wound healing assay using livecell microscopy. Cell Adhes Migr 2014;8:440–51. 10.4161/cam.36224.

76 Achterberg VF, Buscemi L, Diekmann H, Smith-Clerc J, Schwengler H, Meister JJ, et al. The nano-scale mechanical properties of the extracellular matrix regulate dermal fibroblast function. J Invest Dermatol 2014;134:1862–72. 10.1038/jid.2014.90.

77 Pailler-Mattei C, Bec S, Zahouani H. In vivo measurements of the elastic mechanical properties of human skin by indentation tests. Med Eng Phys 2008;30:599–606. 10.1016/j.medengphy.2007.06.011.

78 Kao AP, Connelly JT, Barber AH. 3D nanomechanical evaluations of dermal structures in skin. J Mech Behav Biomed Mater 2016;57:14–23. 10.1016/j.jmbbm.2015.11.017.

79 Chantre CO, Campbell PH, Golecki HM, Buganza AT, Capulli AK, Deravi LF, et al. Production-scale fibronectin nanofibers promote wound closure and tissue repair in a dermal mouse model. Biomaterials 2018;166:96–108. 10.1016/j.biomaterials.2018.03.006.

80 Larsen M, Artym V V., Green JA, Yamada KM. The matrix reorganized: extracellular matrix remodeling and integrin signaling. Curr Opin Cell Biol 2006;18:463–71. 10.1016/j.ceb.2006.08.009.

81 Pussell BA, Peake PW, Brown MA, Charlesworth JA. Human fibronectin metabolism. J Clin Invest 1985;76:143–8. 10.1172/JCI111937.

82 Sherman L, Lee J. Fibronectin: blood turnover in normal animals and during intravascular coagulation. Blood 1982;60:558–63. 10.1182/blood.v60.3.558.558.

83 Deno D, Saba T, Lewis E. Kinetics of endogenously labeled plasma fibronectin: incorporation into tissues. Am J Physiol Integr Comp Physiol 1983;245:R564–75.

84 Kriegsmann J, Berndt A, Hansen T, Borsi L, Zardi L, Bräuer R, et al. Expression of fibronectin splice variants and oncofetal glycosylated fibronectin in the synovial membranes of patients with rheumatoid arthritis and osteoarthritis. Rheumatol Int 2004;24:25–33. 10.1007/s00296-003-0316-1.

85 Shi F, Sottile J. MT1-MMP regulates the turnover and endocytosis of extracellular matrix fibronectin. J Cell Sci 2011;124:4039–50. 10.1242/jcs.087858.

86 Yamada KM, Doyle AD, Lu J. Cell–3D matrix interactions: recent advances and opportunities. Trends Cell Biol 2022;32:883–95. 10.1016/j.tcb.2022.03.002.

87 Das SL, Bose P, Lejeune E, Reich DH, Chen C, Eyckmans J. Extracellular Matrix Alignment Directs Provisional Matrix Assembly and Three Dimensional Fibrous Tissue Closure. Tissue Enigneering 2021;27:. 10.1089/ten.tea.2020.0332.

88 Früh SM, Schoen I, Ries J, Vogel V. Molecular architecture of native fibronectin fibrils. Nat Commun 2015;6:. 10.1038/ncomms8275.

89 Coussen F, Choquet D, Sheetz MP, Erickson HP. Trimers of the fivronectin cell adhesion domain localize to actin filament bundles and undergo rearward translocation. J Cell Sci 2002;115:2581–90. 10.1242/jcs.115.12.2581.

90 Tomer D, Arriagada C, Munshi S, Alexander BE, French B, Vedula P, et al. A new mechanism of fibronectin fibril assembly revealed by live imaging and super-resolution microscopy. J Cell Sci 2022;135:. 10.1242/jcs.260120.

91 Seo BR, Chen X, Ling L, Song YH, Shimpi AA, Choi S, et al. Collagen microarchitecture mechanically controls myofibroblast differentiation. Proc Natl Acad Sci 2020;117:201919394. 10.1073/pnas.1919394117.

92 Elbediwy A, Vincent-Mistiaen ZI, Spencer-Dene B, Stone KR, Boeing S, Wculek SK, et al. Integrin signalling regulates YAP and TAZ to control skin homeostasis. Development 2016:1674.

93 Elosegui-Artola A, Andreu I, Beedle AEM, Lezamiz A, Uroz M, Kosmalska AJ, et al. Force Triggers YAP Nuclear Entry by Regulating Transport across Nuclear Pores. Cell 2017;171:1397–1410.e14. 10.1016/j.cell.2017.10.008.

94 Porazinski S, Wang H, Asaoka Y, Behrndt M, Miyamoto T, Morita H, et al. YAP is essential for tissue tension to ensure vertebrate 3D body shape. Nature 2015;521:217–21. 10.1038/nature14215.

95 Scott KE, Fraley SI, Rangamani P. A spatial model of YAP/TAZ signaling reveals how stiffness, dimensionality, and shape contribute to emergent outcomes. Proc Natl Acad Sci U S A 2021;118:1–12. 10.1073/pnas.2021571118.

96 Shi L, Nadjar-Boger E, Jafarinia H, Carlier A, Wolfenson H. YAP mediates apoptosis through failed integrin adhesion reinforcement. Cell Rep 2024;43:. 10.1016/j.celrep.2024.113811.

97 Goffin JM, Pittet P, Csucs G, Lussi JW, Meister JJ, Hinz B. Focal adhesion size controls tension-dependent recruitment of α-smooth muscle actin to stress fibers. J Cell Biol 2006;172:259–68. 10.1083/jcb.200506179.

98 Dugina V, Fontao L, Chaponnier C, Vasiliev J, Gabbiani G. Focal adhesion features during myofibroblastic differentiation are controlled by intracellular and extracellular factors. J Cell Sci 2001;114:3285–96. 10.1242/jcs.114.18.3285.

99 Cao X, Ban E, Baker BM, Lin Y, Burdick JA, Chen CS, et al. Multiscale model predicts increasing focal adhesion size with decreasing stiffness in fibrous matrices. Proc Natl Acad Sci U S A 2017;114:E4549–55. 10.1073/pnas.1620486114.

100 Engler A, Bacakova L, Newman C, Hategan A, Griffin M, Discher D. Substrate Compliance versus Ligand Density in Cell on Gel Responses. Biophys J 2004;86:617–28. 10.1016/S0006-3495(04)74140-5.

101 Changede R, Cai H, Wind SJ, Sheetz MP. Integrin nanoclusters can bridge thin matrix fibres to form cell–matrix adhesions. Nat Mater 2019;18:1366–75. 10.1038/s41563-019-0460-y.

102 Liu Z, Fu J, Yuan H, Ma B, Cao Z, Chen Y, et al. Polyisocyanide hydrogels with tunable nonlinear elasticity mediate liver carcinoma cell functional response. Acta Biomater 2022;148:152–62. 10.1016/j.actbio.2022.06.022.

103 Brusatin G, Panciera T, Gandin A, Citron A, Piccolo S. Biomaterials and engineered microenvironments to control YAP/TAZ-dependent cell behaviour. Nat Mater 2018;17:1063–75. 10.1038/s41563-018-0180-8.

104 Walker M, Pringle EW, Ciccone G, Oliver-Cervelló L, Tassieri M, Gourdon D, et al. Mind the Viscous Modulus: The Mechanotransductive Response to the Viscous Nature of Isoelastic Matrices Regulates Stem Cell Chondrogenesis. Adv Healthc Mater 2023;2302571:1–17. 10.1002/adhm.202302571.

105 Raghunathan VK, Morgan JT, Dreier B, Reilly CM, Thomasy SM, Wood JA, et al. Role of substratum stiffness in modulating genes associated with extracellular matrix and mechanotransducers YAP and TAZ. Investig Ophthalmol Vis Sci 2013;54:378–86. 10.1167/iovs.12-11007.

106 Mugahid D, Kalocsay M, Liu X, Gruver JS, Peshkin L, Kirschner MW. YAP regulates cell size and growth dynamics via non-cell autonomous mediators. Elife 2020;9:1–20. 10.7554/eLife.53404.

107 Pocaterra A, Romani P, Dupont S. YAP/TAZ functions and their regulation at a glance. J Cell Sci 2020;133:1–9. 10.1242/jcs.230425.

108 Sero JE, Bakal C. Multiparametric Analysis of Cell Shape Demonstrates that β-PIX Directly Couples YAP Activation to Extracellular Matrix Adhesion. Cell Syst 2017;4:84–96.e6. 10.1016/j.cels.2016.11.015.

109 Farhadifar R, Röper JC, Aigouy B, Eaton S, Jülicher F. The Influence of Cell Mechanics, Cell-Cell Interactions, and Proliferation on Epithelial Packing. Curr Biol 2007;17:2095–104. 10.1016/j.cub.2007.11.049.

110 Rognoni E, Pisco AO, Hiratsuka T, Sipilä KH, Belmonte JM, Mobasseri SA, et al. Fibroblast state switching orchestrates dermal maturation and wound healing. Mol Syst Biol 2018;14:1–20. 10.15252/msb.20178174.

111 Conway JRW, Isomursu A, Follian G, Harma V, Jou-Olle E, Pasquier N, et al. Defined extracellular matrix compositions support stiffness-insensitive cell spreading and adhesion signaling James. Proc Natl Acad Sci 2017;120:. 10.1073/pnas.

112 Nardone G, Oliver-De La Cruz J, Vrbsky J, Martini C, Pribyl J, Skládal P, et al. YAP regulates cell mechanics by controlling focal adhesion assembly. Nat Commun 2017;8:. 10.1038/ncomms15321.

113 Venghateri JB, Dassa B, Morgenstern D, Shreberk-Shaked M, Oren M, Geiger B. Deciphering the involvement of the Hippo pathway co-regulators, YAP/TAZ in invadopodia formation and matrix degradation. Cell Death Dis 2023;14:. 10.1038/s41419-023-05769-1.

114 An Seong SA, Llinás M, Jimenez-Barbero J, Petersen TE. The Two Polypeptide Chains in Fibronectin Are Joined in Antiparallel Fashion:NMR Structural Characterization. Biochemistry 1992;31:9927–33. 10.1021/bi00156a010.

115 Arnoldini S, Moscaroli A, Chabria M, Hilbert M, Hertig S, Schibli R, et al. Novel peptide probes to assess the tensional state of fibronectin fibers in cancer. Nat Commun 2017;8:. 10.1038/s41467-017-01846-0.

116 Chantre CO, Campbell PH, Golecki HM, Buganza AT, Capulli AK, Deravi LF, et al. Production-scale fibronectin nanofibers promote wound closure and tissue repair in a dermal mouse model. Biomaterials 2018;166:96–108. 10.1016/j.biomaterials.2018.03.006.

